# Evaluating the potential of cross-species neutralization of anti-PfCyRPA and anti-PfRIPR Monoclonal Antibodies

**DOI:** 10.1101/2025.11.03.682341

**Authors:** Noemi Guerra, Kelly A. Hagadorn, Megha Nair, Laty Gaye Thiam, Barnabas G. Williams, Kirsty McHugh, Dimitra Pipini, Julie Healer, Paul Masendycz, Alan F. Cowman, Simon J. Draper, Amy K. Bei

## Abstract

PfCyRPA and PfRIPR are promising next-generation malaria blood-stage vaccine candidate antigens that play an essential role in erythrocyte invasion of *Plasmodium falciparum*. CyRPA and RIPR orthologs are present in all human-infecting *Plasmodium* species, suggesting the potential for a cross-species vaccine. Using Growth Inhibition Assays (GIA), this study investigates seven anti-PfCyRPA and three anti-PfRIPR monoclonal antibodies targeting *P. falciparum* for their inhibitory activity against *P. knowlesi*, a non-falciparum species that contributes to a significant burden of zoonotic disease in South-East Asia, shares some biological features with *Plasmodium vivax*, and has a robust *in vitro* culture system. Despite their efficacy against *P. falciparum* and partially conserved epitopes, these antibodies exhibited minimal inhibition of *P. knowlesi*. Understanding the antigenic diversity and immune mechanisms across *Plasmodium* species is critical for advancing pan-species vaccine strategies.

## Background

Malaria remains a major global health concern. In 2023 an estimated 263 million cases and 597,000 deaths attributed to malaria globally. While there are six human-infecting *Plasmodium* species, P. falciparum has been the main target of global health interventions due to its significant contribution to morbidity and mortality. Currently, the WHO recommends two vaccines, RTS,S/AS01 and R21/Matrix-M, for children living in areas with moderate to high *P. falciparum* transmission, both of which target the pre-erythrocytic stage of *P. falciparum*. Next-generation vaccines targeting other life-cycle stages are critical to explore, either independently or in combination, with the goal of achieving a highly efficacious strain-transcendent malaria vaccine to further reduce disease burden. Promising blood-stage vaccine candidate antigens currently being pursued include PTRAMP, CSS, RIPR, CyRPA, and RH5, which together form the PCRCR pentameric protein complex of *P. falciparum*, and all of which are conserved and essential for merozoite invasion of erythrocytes^1^. Of the PCRCR-complex, vaccines targeting PfRH5 are the furthest along in clinical development, with RH5.1/Matrix-M in a Phase 2b field efficacy trial (ClinicalTrials.gov NCT05790889)^2^. Combination vaccines incorporating RH5 and other members of the PCRCR-complex, such as CyRPA and RIPR, have also entered Phase 1 clinical trials with the goal of enhancing the efficacy of the RH5.1 vaccine (ClinicalTrials.gov NCT05385471).

Additional potential benefits of vaccines targeting other members of the PCRCR-complex include vaccines that could target multiple Plasmodium species. While RH5 is only found in P. falciparum and in the Laverania subgenus, other members of the PCRCR-complex such as CyRPA and RIPR are present in most *Plasmodium* species^3^. After *P. falciparum, P. vivax* presents the greatest global health burden, responsible for 48.2% of malaria cases in South-East Asia and 72.1% in the Americas. Currently, no vaccine targeting multiple *Plasmodium* species exists, which is partially due to the lack of in vitro culture systems for *P. vivax, P. malariae, P. ovale curtisi*, and *P. ovale wallikeri* making *in vitro* experimental approaches more challenging. Of the two species with robust *in vitro* culture systems (*P. falciparum* and *P. knowlesi*), *P. knowlesi* is most genetically similar to *P. vivax, P. malariae, P. ovale curtisi*, and *P. ovale wallikeri*^4^. *P. knowlesi* is a zoonotic malaria that mainly circulates in macaques but regularly infects humans. The biology of *P. knowlesi* is very similar to that of *P. vivax* in that it is dependent on the DARC receptor for red blood cell invasion while lacking the reticulocyte restriction characteristic of *P. vivax*.

The goal of this study was to evaluate whether anti-PfCyRPA and anti-PfRIPR mAbs, previously shown to possess invasion inhibitory properties against *P. falciparum* laboratory strains, could similarly inhibit *P. knowlesi*. We thus sought to determine whether CyRPA and RIPR orthologues are similar enough to permit the development of not only a strain-transcendent vaccine within a species, but potentially a species-transcendent vaccine across non-falciparum Plasmodium species. Specifically, we aimed to characterize inhibitory profiles of anti-PfCyRPA and anti-PfRIPR mAbs against laboratory lines of *P. falciparum* and *P. knowlesi*.

## Materials and Methods

### Amino Acid Alignments and Epitope Mapping

*P. falciparum* and *P. knowlesi* CyRPA (PF3D7_0423800 - 812432 and PKNH_0515800 – 7319417) and RIPR (PF3D7_0323400 – 814549 and PKNH_0817000 - 7320243) genes were aligned in Geneious Prime 2025.0.3 using MUSCLE 5.1. Antibody binding epitopes were previously described^5–8^.

### Monoclonal Antibody Preparation

Seven previously characterized anti-PfCyRPA mAbs (Cy.002, Cy.003, Cy.004, Cy.007, Cy.009 [5], 8A7, and 5B12 [7]) and three previously characterized anti-PfRIPR mAbs (1C4, 5G6, 1G12 [6]) were produced as previously described^5–8^). Briefly, Cy.003, Cy.004, Cy.007, and Cy.009 mAbs were produced by Icosagen using HybriFree Technology. Spleen cells from immunized chickens were panned against antigen-coated immune modules. RNA that was extracted from bound cells were used to synthesize cDNA and amplify variable light and heavy chains. Amplified variable light and heavy chains were purified and cloned into human immunoglobulin G1 expression vectors^5^ Cy.002, 8A7, and 5B12 were produced from immunized mice, while 1C4, 5G6, and 1G12 were produced from immunized rabbits. Spleen cells from immunized mice or rabbits were fused with myeloma cells and positive antibody-producing hybridoma cell lines were identified using ELISA. For 8A7, 5B12, 1C4, 5G6, and 1G12, positive hybridomas were sub-cloned to obtain monoclonal cell lines^6,7^. For Cy.002 and 8A7, variable domains of the heavy and light chains were sequenced, synthesized, and cloned into AbVec-hIgG1/AbVec-hIgG1-kappa vectors^5^. Anti-PfCyRPA and -PfRIPR mAbs were buffer exchanged in unsupplemented RPMI culture media using 30kDa-cut-off centrifugal filter units (EMD Millipore) and resuspended in unsupplemented RMPI culture media at 1 mg/ml.

### Parasite Culture

3D7 *P. falciparum* cultures were maintained at 0.5-1.5% parasitemia and 4% hematocrit in supplemented parasite medium (RPMI 1640 containing 25 mM HEPES, 0.1 mg/mL Hypoxanthine, and 50*µ*g/mL Gentamicin and supplemented with Albumax II (0.5% v/v), 2 mg/mL sodium bicarbonate, and heat-inactivated human AB serum (5% v/v). H1 *P. knowlesi* cultures (human-RBC adapted clone YH1) were maintained at 0.5-1.5% parasitemia and 4% hematocrit in supplemented parasite medium (RPMI 1640 containing 25 mM HEPES, 0.1mg/ml Hypoxanthine, and 50 *µ*g/mL Gentamicin and supplemented with Albumax II (0.5% v/v), 0.292 mg/mL of L-glutamine, 2 mg/mL sodium bicarbonate, and horse serum (10% v/v)). Human erythrocytes of blood group O+ donors, who were also Duffy positive were used for all cultures and cultures were incubated in an atmosphere of 1% O2, 5% CO2, and 94% N2 at 37 °C.

### Growth Inhibition Assays

Growth inhibition assays (GIAs) quantify the anti-merozoite inhibitory activity of antibodies. Prior to setting up GIAs, H1 *P. knowlesi* cultures were synchronized with a 280mM D-sorbitol, 20mM Na-HEPES, and 0.1 mg/ml BSA solution (“*P. knowlesi* sorbitol”), while 3D7 *P. falciparum* cultures were synchronized with 5% D-sorbitol to kill non-ring blood stages. For GIAs, mAbs were added to 1% parasitemia at 4% hematocrit cultures in 96 well half-well area plates (for a final concentration of 1% parasitemia and 2% hematocrit) with a total volume of 40*µ*L and incubated for one full erythrocytic cycle (approximately 24h for *P. knowlesi* H1 and approximately 48h for *P. falciparum* 3D7). Anti-PfCyRPA mAbs were tested in duplicate at final antibody concentrations of 200*µ*g/ml, 100*µ*g/ml, 50*µ*g/ml, and 25*µ*g/ml. Three biological replicates were conducted. To evaluate synergistic antibody effect, determining if combing mAbs resulted in a greater inhibition effect than if the mAbs were evaluated alone^8^, combinations of anti-PfCyRPA mAb were performed^5,9^. Anti-PfCyRPA mAbs used in combination (Cy.003 : Cy.009, Cy.004 : Cy.007, Cy.007 : Cy.009) were combined at equal concentrations (1 : 1) to achieve the same final antibody concentrations previously mentioned (i.e. Cy.003 100*µ*g/ml + Cy.009 100*µ*g/ml = Cy.003 : Cy.009 200*µ*g/ml). Anti-PfRIPR mAbs were tested in duplicate at concentrations of 400*µ*g/ml, 200*µ*g/ml, 100*µ*g/ml, and 50*µ*g/ml). Two biological replicates were conducted. For controls, a-BSG (Anti-CD147 antibody [MEM-M6/6], abcam, ab119114) was tested at 10*µ*g/ml, 1*µ*g/ml, and 0.1*µ*g/ml and anti-DARC (DARC Antibody (2C3), Novus Biologicals, NBP2-75196-0.025mg) was tested at 10*µ*g/ml, 1*µ*g/ml, and 0.1*µ*g/ml as well as 200*µ*g/ml, 100 *µ*g/ml, 50 *µ*g/ml, and 25*µ*g/ml for two of the H1 *P. knowlesi* and 3D7 *P. falciparum* biological triplicate experiments. Naive total IgG was tested at all the listed concentrations. After at least 95% of the parasites re-invaded in control wells with no antibody (RPMI only), the assays were harvested. Parasites cultures were transferred to 96 well U-bottom plates, washed once with 1X PBS-1% BSA, stained with a dilution of 1/2000 SYBR Green I (10,000X stock, Invitrogen) in PBS for 20 min, washed twice with 1X PBS-1% BSA, and resuspended in 1X PBS^10^. Parasitemia for each well was acquired using a Cytoflex cytometer (Beckman Coulter), with 100,000 events (erythrocytes) per sample. Flow cytometry data were analyzed using Flow Jo_v10.8.1 software. Percent inhibition was calculated as 100-[(Parasitemia of Test Antibody)/(Parasitemia of Naïve IgG control)x100].

## Results

### *P. falciparum* and *P. knowlesi* CyRPA and RIPR Amino Acid Alignment and mAb epitope mapping

*P. falciparum* and *P. knowlesi* genes aligned with 21.3% identity for CyRPA and 44.6% identity for RIPR using MUSCLE 5.1 (Figure 1). The contact residues between PfCyRPA mAbs Cy.002, Cy.003, Cy.004, Cy.007, 8A7, and PfCyRPA can be determined from their crystal structure^5^. A high-resolution structure of Cy.009 is not available; however, it is known to compete with Cy.004 for binding^5^. Similarly, mapped contact residues between 5B12 and PfCyRPA have not been identified, but was found to compete for binding with 8A7^6^. For anti-PfRIPR mAbs 1C4, 5G6, 1G12, binding to PfRIPR has only been identified to specific binding regions (i.e EGF-like domain specific)^6^(Figure 1B).

**Figure 1.**
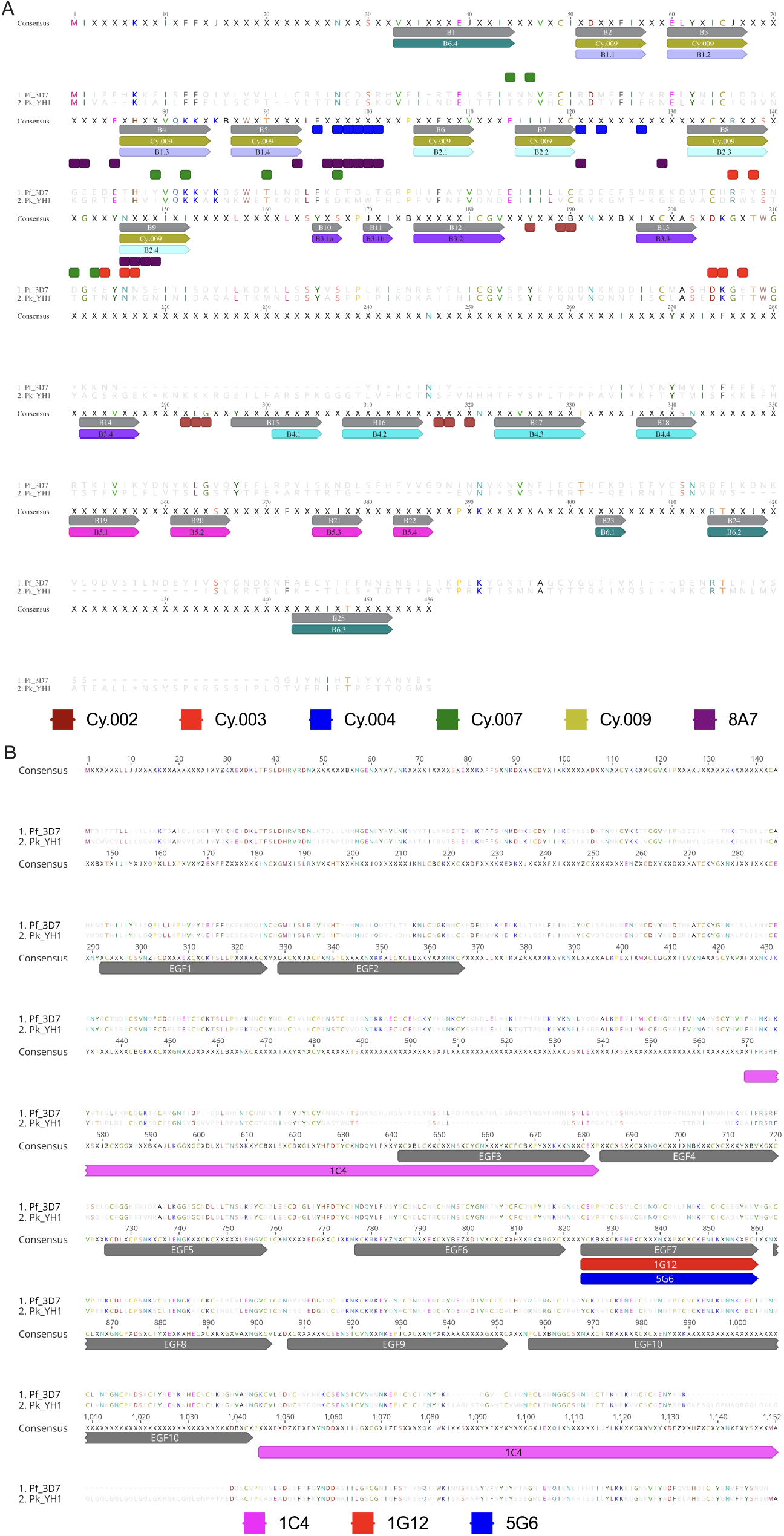
*P. falciparum* and *P. knowlesi* CyRPA and RIPR Gene Alignments. A. Aligned *P. falciparum* and *P. knowlesi* CyRPA genes. Alignment is annotated with contact residues between PfCyRPA mAbs and PfCyRPA (Cy.002 - dark red, Cy.003 - red, Cy.004 - blue, Cy.007 - green, and 8A7 - purple) and with PfCyRPA mAbs to specific PfCyRPA binding regions when contact residues are unknown (i.e blade specific – light blue and light purple) (Cy.009 - yellow). CyRPA blades are annotated as Blade number.beta-sheet (e.g. - Blade 1, beta-sheet 2 = B1.2). B. Aligned *P. falciparum* and *P. knowlesi* RIPR genes. Alignment is annotated with PfRIPR mAbs to specific PfRIPR binding regions when contact residues are unknown (i.e EGF domain specific - gray) (1C4 - pink, 5G6 - red, 1G12 - blue).

### Anti-PfCyRPA monoclonal antibodies inhibit *P. falciparum*, but not *P. knowlesi* in GIA

Seven of the anti-PfCyRPA mAbs tested reduced growth of *P. falciparum*, with a range of 13.4% to 69.8% inhibition at the highest mAb concentration (Figure 2A). When mAbs were tested individually against Pf3D7, Cy.009 demonstrated the most potent growth inhibition response, with inhibition reaching 69.8% at 200*µ*g/ml. The highest growth inhibition response achieved against Pf3D7 was when anti-PfCyRPA mAbs were used in combinations, reaching inhibition up to 77.9% for Cy.007 : Cy.009 at 200 *µ*g/ml. Against Pf3D7, synergy was observed for all combinations (Cy.003 : Cy.009, Cy.004 : Cy.007, Cy.007 : Cy.009) at 100*µ*g/ml, 50*µ*g/ml, and 25*µ*g/ml, consistent with previous studies^5^. However, the same mAbs demonstrated no growth inhibition against *P. knowlesi* (−7.5% to −0.37%) even when used in known synergistic combinations (Figure 2B). Positive controls included an anti-DARC mAb for *P. knowlesi*, as *P. knowlesi* relies on DARC as an invasion receptor^11^, and an anti-BSG mAb for *P. falciparum*, as *P. falciparum* relies on basigin as an invasion receptor^12^. The anti-DARC mAb reached an inhibition of 88.2% against *P. knowlesi*, while anti-BSG mAb showed no inhibition (Figure 2B).

**Figure 2.**
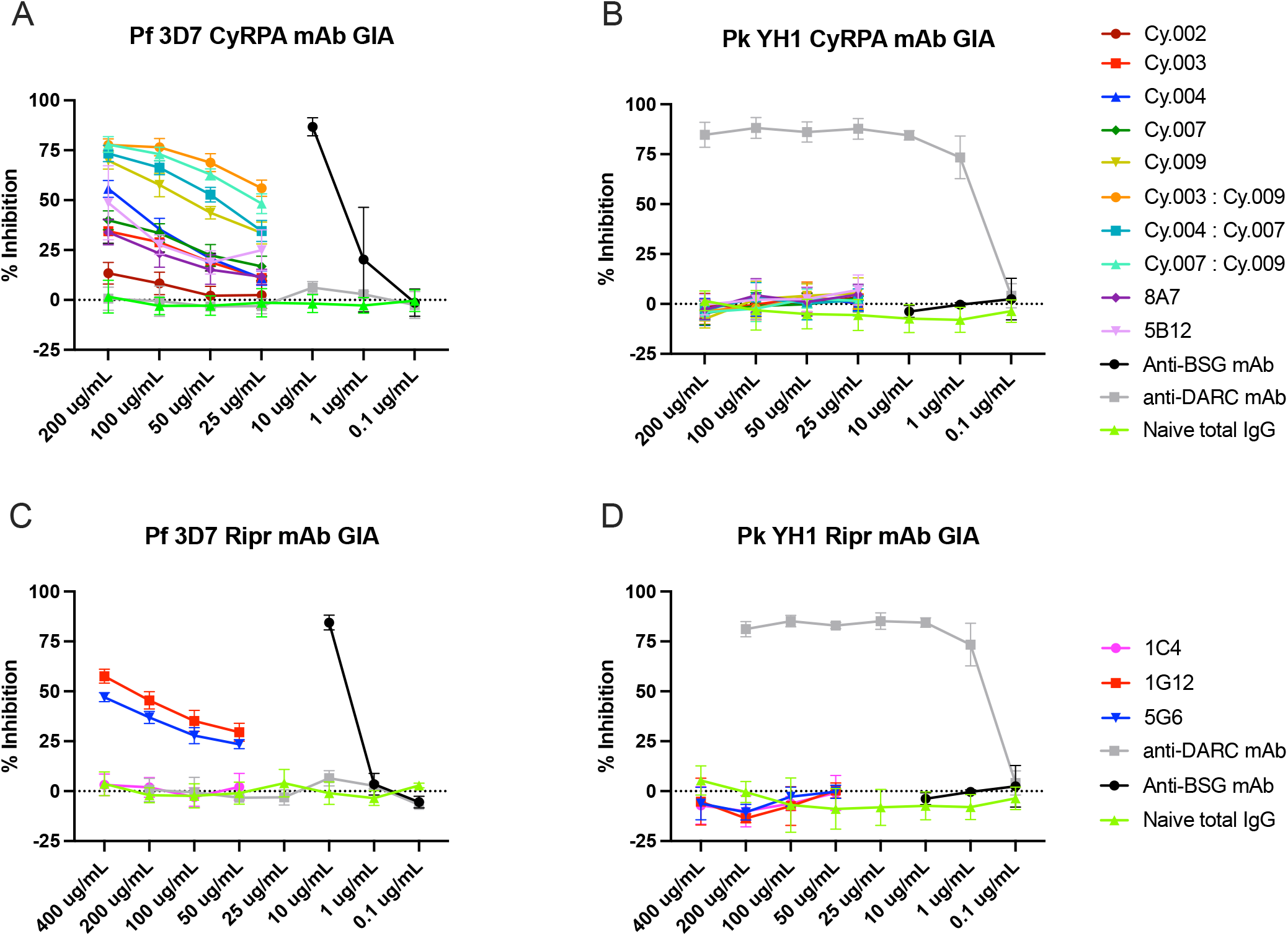
Assessment of anti-PfCyRPA and anti-PfRIPR monoclonal antibodies against 3D7 *P. falciparum* and H1 *P. knowlesi*. A. Growth inhibitory activity of seven anti-PfCyRPA mAbs, alone or in combination, tested against 3D7 *P. falciparum*. B. Growth inhibitory activity of seven anti-PfCyRPA mAbs, alone or in combination, tested against YH1 *P. knowlesi*. C. Growth inhibitory activity of three anti-PfRIPR mAbs tested at different concentration against 3D7 *P. falciparum*. D. Growth inhibitory activity of three anti-PfRIPR mAbs tested at different concentration against YH1 *P. knowlesi*. Controls include Naïve total IgG, anti-DARC mAb, and anti-BSG mAb. Anti-DARC and anti-BSG mAbs serve as positive and negative controls dependent on *Plasmodium* species. All experiments were performed with technical duplicates. N = 3 biological replicates were performed for anti-PfCyRPA mAbs and N = 2 biological replicates were performed for anti-PfRIPR mAbs. Error bars show SD.

### Anti-PfRIPR monoclonal antibodies inhibit *P. falciparum*, but not *P. knowlesi* in GIA

Similar to anti-PfCyRPA mAbs, the three anti-PfRIPR mAbs reduced growth against *P. falciparum* and ranged from 3.2% to 57.6% at the highest mAb concentration (Figure 2C). Against Pf3D7, 1G12 had the most potent growth inhibition response, with inhibition reaching 57.6% at 400 *µ*g/ml. 1C4 had no growth inhibition response against Pf3D7, only achieving 3.2% at 400*µ*g/ml, similar to the observed Naive total IgG response (3.7% at 400*µ*g/ml). Again, no growth inhibition was demonstrated against *P. knowlesi* (−7.1% to −5.2%) despite anti-*P. falciparum* activity (Figure 2D). The anti-BSG mAb reached an inhibition of 84.5% against *P. falciparum*, while the anti-DARC mAb showed no inhibition (Figure 2D).

## Discussion

With the bold goal of developing a species-transcendent vaccine across non-falciparum *Plasmodium* species, CyRPA and RIPR orthologs are attractive candidates as they exist in all human-infecting Plasmodium species. Here, our goal was to evaluate the potential for cross-species antibody-mediated inhibition of antibodies targeting CyRPA and RIPR. While these seven anti-PfCyRPA mAbs and the three anti-PfRIPR mAbs have been tested against *P. falciparum* previously and demonstrated a range of invasion inhibition, their potential for cross-species antibody-mediated inhibition had not been evaluated. Although amino acid identity overall for *P. falciparum* and *P. knowlesi* was low for CyRPA (21.3%) and RIPR (44.6%), identity was as high as high as 50% for amino acids within known mAb binding epitopes (Cy.002, Cy.003, Cy.004, Cy.007, Cy.009, 8A7) and 64% for mAbs with unknown specific epitopes but known domain binding regions (1C4, 1G12, 5G6). For 5B12, mapped contact residues to PfCyRPA are not known. Our rationale for testing these anti-PfCyRPA and anti-PfRIPR mAbs against *P. knowlesi* was that some binding epitopes may be shared between *P. falciparum* and *P. knowlesi*, some epitopes (e.g., 5B12) remain undefined and could be conserved, and cross-species inhibition against *P. knowlesi* has previously been observed with antigen-specific antibodies^13,14^.

While the specific mAbs tested did not display any cross-species inhibition, these findings remain informative, highlighting epitope differences and functional differences in CyRPA and RIPR between *P. falciparum* vs *P. knowlesi*. Our anti-DARC mAb and anti-BSG mAb controls displayed inhibition as expected for their respective invasion receptor in *P. falciparum* and *P. knowlesi*, consistent with previous studies^3^. In *P. knowlesi*, it is reported that PkRIPR together with PkPTRAMP and PkCSS form a conserved invasion scaffold, but the erythrocyte binding proteins and receptors are currently unknown^3,15^. PkCyRPA is essential for parasite growth but is not believed to be part of the PCR-complex in *P. knowlesi*^3^. Given these differences between CyRPA and RIPR in *P. falciparum* and *P. knowlesi*, the reason for the lack of inhibition observed in our study maybe be due to a multitude of factors including low identity in mAb binding epitopes, epitope sites which could be structurally hidden in one species while exposed in another, and/or functional differences between Plasmodium species during erythrocyte invasion.

While inhibition of *P. knowlesi* has been observed from anti-PkRIPR IgG purified from rabbits immunized with recombinant PkRIPR proteins^3^, a lack of inhibition of *P. knowlesi* was demonstrated from anti-PvCyRPA IgG purified from rabbits immunized with a recombinant PvCyRPA protein^14^. However, excitingly, a recent study showed that PTRAMP, CSS, and RIPR monoclonal antibodies (mAb) and nanobodies generated against *P. vivax* displayed cross-species inhibition against *P. falciparum* and *P. knowlesi*^15^. Evidence of cross-species inhibition and the *in vitro* culture system of *P. knowlesi* warrants further investigation of the cross-species inhibition potential of additional mAbs to conserved invasion complexes.

Much work has been done on the identification of strain-transcendent antibodies against *P. falciparum*, and this represents an important challenge in malaria vaccinology. Here, we argue that concurrent work should be dedicated to the identification of species-transcendent antibodies against human-infecting *Plasmodium* species. The presence of PTRAMP, CSS, CyRPA, and RIPR orthologs in human-infecting *Plasmodium* species make them attractive targets. Future studies should explore the cross-strain and cross-species activity of antibodies isolated from human clinical trials as the mAbs tested here were isolated from animals. To enhance the likelihood of cross-species inhibition, it will be critical to identify epitopes that are maximally conserved across orthologs through detailed epitope mapping. In conclusion, next-generation vaccine candidates present the intriguing possibility of targeting all human-infecting malaria species; however, to maximize this potential, it is important to consider the functional differences that exist between Plasmodium species, to incorporate detailed epitope mapping of mAbs, and to employ highly sensitive and rigorous phenotypic screening such as the GIA assay described here.

## Acknowledgments

YH1 was kindly provided by Manoj T. Duraisingh, *P. knowlesi* culture protocols were kindly provided by Robert A. Moon. Standardized IgG purification protocol for GIA was kindly provided by Carole A. Long. EURIPRED generated mAbs, Cy.003 (catalogued as 3B3#17), Cy.004 (4D12#30), Cy.007 (3A7#22) and Cy.009 (7B9#13) are available only through the National Institute of Biological Standards and Control, UK. These mAbs were produced through the European Commission FP7 EURIPRED project (INFRA-2012-312661), funded by the European Union’s Seventh Framework Programme [FP7/2007—2013] under Grant Agreement No: 312661—European Research Infrastructures for Poverty Related Diseases (EURIPRED).

## Funding

LGT is supported by an ARISE grant from the African Academy of Sciences (ARISE-PP-FA-056). KAH is supported by National Institute of Allergy and Infectious Diseases of the NIH (1F31AI176818-01A1) and the Yale Center for Clinical Investigation (TR001864). NG is supported by a Yale STARS Fellowship. The work is supported by the National Institute of Allergy and Infectious Diseases of the NIH (R01 AI168238) to AKB. Cy.003, Cy.004, Cy.007, and Cy.009 were provided by Icosagen AS through the Centre for AIDS Reagents repository at the National Institute for Biological Standards and Control, UK. These mAbs were produced through the European Commission FP7 EURIPRED project (INFRA-2012-312661), funded by the European Union’s Seventh Framework Programme [FP7/2007—2013] under Grant Agreement No: 312661—European Research Infrastructures for Poverty Related Diseases (EURIPRED).

## Data Availability

GIA data have been deposited in the Dryad database and is publicly available at: https://doi.org/10.5061/dryad.0cfxpnwfw.

## Author contributions

Project conception, management and coordination: N.G., K.A.H., A.K.B. Data collection and experimentation: N.G., K.A.H., M.N., A.K.B. Data analysis: N.G., K.A.H., M.N., A.K.B. Contribution of reagents, materials, and analysis tools: L.G.T., B.G.W., K.M., D.P., J.H., P.M., A.F.C., S.J.D., A.K.B. Paper writing: N.G., K.A.H., A.K.B. All authors read and approved the manuscript.

## Competing interests

SJD, BGW, KM are inventors on patent applications relating to RH5 and/or RCR-complex malaria vaccines and/or antibodies. The other authors declare no conflicts of interest.

## Presentation of the presented findings

Portions of this work were presented at the Multilateral Initiative on Malaria Meeting in 2024. Guerra N, Ba A, Pouye MN, Thiam LG, Hagadorn K, Nair M, McHugh K, Pipini D, Sene SD, Wade A, Mbengue A, Draper SJ, Bei AK. Investigating Cross-Strain and Cross-Species Neutralization of Anti-CyRPA Antibodies in Clinical Isolates and Chimeras. Multilateral Initiative on Malaria (MIM) Meeting, Kigali, Rwanda. 2024.

## Notes

https://doi.org/10.5061/dryad.0cfxpnwfw

## References

1. Scally, S. W. et al. Pcrcr complex is essential for invasion of human erythrocytes by plasmodium falciparum. Nat Microbiol 7, 2039–2053 (2022).

2. Natama, H. M. et al. Safety and efficacy of the blood-stage malaria vaccine rh5.1/matrix-m in burkina faso: interim results of a double-blind, randomised, controlled, phase 2b trial in children. Lancet Infect Dis 25, 495–506 (2025).

3. Knuepfer, E. et al. Divergent roles for the rh5 complex components, cyrpa and ripr in human-infective malaria parasites. PLoS Pathog 15, e1007809 (2019).

4. Rutledge, G. G. et al. Plasmodium malariae and p. ovale genomes provide insights into malaria parasite evolution. Nature 542, 101–104 (2017).

5. Ragotte, R. J. et al. Heterotypic interactions drive antibody synergy against a malaria vaccine candidate. Nat Commun 13, 933 (2022).

6. Healer, J. et al. Neutralising antibodies block the function of rh5/ripr/cyrpa complex during invasion of plasmodium falciparum into human erythrocytes. Cell Microbiol 21, e13030 (2019).

7. Chen, L. et al. Structural basis for inhibition of erythrocyte invasion by antibodies to plasmodium falciparum protein cyrpa. Elife 6 (2017).

8. Knudsen, A. S. et al. Strain-dependent inhibition of erythrocyte invasion by monoclonal antibodies against plasmodium falciparum cyrpa. Front Immunol 12, 716305 (2021).

9. Azasi, Y. et al. Bliss’ and loewe’s additive and synergistic effects in plasmodium falciparum growth inhibition by ama1-ron2l, rh5, ripr and cyrpa antibody combinations. Sci Rep 10, 11802 (2020).

10. Bei, A. K. & Duraisingh, M. T. Measuring plasmodium falciparum erythrocyte invasion phenotypes using flow cytometry. Methods Mol Biol 1325, 167–86 (2015).

11. Miller, L. H., Mason, S. J., Dvorak, J. A., McGinniss, M. H. & Rothman, I. K. Erythrocyte receptors for (plasmodium knowlesi) malaria: Duffy blood group determinants. Science 189, 561–3 (1975).

12. Crosnier, C. et al. Basigin is a receptor essential for erythrocyte invasion by plasmodium falciparum. Nature 480, 534–7 (2011).

13. Muh, F. et al. Cross-species reactivity of antibodies against plasmodium vivax blood-stage antigens to plasmodium knowlesi. PLoS Negl Trop Dis 14, e0008323 (2020).

14. Ndegwa, D. N. et al. Using plasmodium knowlesi as a model for screening plasmodium vivax blood-stage malaria vaccine targets reveals new candidates. PLoS Pathog 17, e1008864 (2021).

15. Seager, B. A. et al. Ptramp, css and ripr form a conserved complex required for merozoite invasion of plasmodium species into erythrocytes. bioRxiv (2025).

